# Effect of β-chitosan on the binding interaction between SARS-CoV-2 S-RBD and ACE2

**DOI:** 10.1101/2020.07.31.229781

**Authors:** Gulimiran Alitongbieke, Xiu-min Li, Qi-Ci Wu, Zhi-Chao Lin, Jia-Fu Huang, Yu Xue, Jing-Na Liu, Jin-Mei Lin, Tao Pan, Yi-Xuan Chen, Yi Su, Guo-Guang Zhang, Bo Leng, Shu-Wen Liu, Yu-Tian Pan

## Abstract

SARS-CoV-2 invades human respiratory epithelial cells via an interaction between its spike RBD protein (SARS-CoV-2 S-RBD) and the host cell receptor angiotensin converting enzyme II (ACE2). Blocking this interaction provides a potent approach to preventing and controlling SARS-CoV-2 infection. In this work, the ability of β-chitosan to block the binding interaction between SARS-CoV-2 S-RBD and ACE2 was investigated. The inhibitory effect of β-chitosan on inflammation induced by the SARS-CoV-2 S-RBD was also studied. Native-PAGE analysis indicated that β-chitosan could bind with ACE2 and the SARS-CoV-2 S-RBD and a conjugate of β-chitosan and ACE2 could no longer bind with the SARS-CoV-2 S-RBD. HPLC analysis suggested that a conjugate of β-chitosan and the SARS-CoV-2 S-RBD displayed high binding affinity without dissociation under high pressure (40 MPa) compared with that of β-chitosan and ACE2. Furthermore, immunofluorescent staining of Vero E6 cells and lungs from hACE2 mice showed that the presence of β-chitosan prevented SARS-CoV-2 S-RBD from binding to ACE2. Meanwhile, β-chitosan could dramatically suppress the inflammation caused by the presence of the SARS-CoV-2 S-RBD both *in vitro* and *vivo*. Moreover, the decreased expression of ACE2 caused by β-chitosan treatment was restored by addition of TAPI-1, an inhibitor of the transmembrane protease ADAM17. Our findings demonstrated that β-chitosan displays an antibody-like function capable of neutralizing the SARS-CoV-2 S-RBD and effectively preventing the binding of the SARS-CoV-2 S-RBD to ACE2. Moreover, ADAM17 activation induced by β-chitosan treatment can enhance the cleavage of the extracellular domain of ACE2, releasing the active ectodomain into the extracellular environment, which can prevent the binding, internalization, and degradation of ACE2 bound to the SARS-CoV-2 S-RBD and thus diminish inflammation. Our study provides an alternative avenue for preventing SARS-CoV-2 infection using β-chitosan.

## INTRODUCTION

According to the latest WHO data (10 September, 2020), the Coronavirus Disease 2019 (COVID-19) pandemic has spread to more than 200 countries and regions and caused more than 27 million confirmed infections, as well as 899 thousand deaths to date, presenting a major threat to public health. Although SARS-CoV-2 was quickly discovered and identified as the pathogen driving the disease outbreak^1,2^, no successful vaccines or specific drugs have been identified and put to use thus far. As a result, the daily numbers of confirmed infections and deaths are still climbing. Therefore, elucidating the mechanisms underlying COVID-19 pathogenesis and developing effective therapeutics to prevent and control the current pandemic have become of the utmost importance.

At the molecular level it has been shown that SARS-CoV-2 (2019-nCoV) uses the same cell entry receptor (ACE2) as SARS-CoV^2^. Bioinformatic analyses also revealed that the spike protein of SARS-CoV-2 is similar to that of SARS-CoV, suggesting that SARS-CoV-2 can infect host cells by binding to ACE2 on the surface of host cells through its spike protein^3^. Hoffmann^4^ then confirmed at the cellular level that SARS-CoV-2 uses the same receptor, ACE2, to enter host cells as SARS-CoV. As a result of these findings, focus shifted to identifying which tissues and organs express ACE2. Several studies reported that ACE2 was not only highly expressed in the lungs, upper esophagus, and stratified squamous epithelium, but also in absorptive enterocytes from the ileum of the small intestine and the colon^5,6^. This led to speculation that SARS-CoV-2 may bind with ACE2 to invade and damage the relevant tissues in infected patients. Since SARS-CoV-2 and SARS-CoV have similar gene sequences, the same binding receptor, and similar clinical manifestations, it is plausible that there may be similar mechanisms of pathogenesis for the two viruses. Endocytosis is induced by the binding of virus to ACE2, resulting in the degradation of the majority of ACE2 and significantly increased angiotensin II (Ang □) levels, which leads to a cytokine storm and multiple organ damage^7^. Structural analysis of ACE2 may help facilitate the design of therapeutic antibodies to prevent SARS-CoV-2 infection. Yan et al. ^8^ were the first to present cryo-electron microscopy structures of full-length human ACE2 and pointed out that the spike protein of SARS-CoV-2 binds to the extracellular peptidase domain of ACE2 mainly through polar residues. Wrapp et al.^9^ obtained a 3.5-angstrom-resolution cryo-electron microscopy structure of the 2019-nCoV spike in the prefusion conformation, and demonstrated that the affinity of ACE2 binding to the 2019-nCoV spike was 10-20 folds greater than the observed affinity for ACE2 binding with the SARS-CoV spike, suggesting a rationale for the higher infectivity and pathogenicity rate of 2019-nCoV compared to SARS-CoV. Thus, it is of great significance to identify molecules that could prevent the binding of SARS-CoV-2 with ACE2 in order to prevent and control the spread of COVID-19.

ACE2 that can protect from severe acute lung failure is the receptor of SARS-CoV-2, which has posed a dilemma for the development of therapies targeted at ACE2. Many reports have confirmed that the ACE2 and ACE genes originated from the same common ancestor^10,11^, making human ACE2 closely related to human ACE. Several studies have confirmed that chitosan binds to the active site of ACE to inhibit its activity, exerting an antihypertensive effect^12,13^. Moreover, since it results in no toxic and side effects for humans, chitosan has become a subject of much research as an antihypertensive functional food and adjuvant drug for various conditions. When comparing the structures of ACE and ACE2 (PDB ID codes 1O8A and 1R42, respectively), it is apparent that both proteins have the same active zinc-binding motif (HEMGH). This implies that ACE2 can likely also bind with chitosan since it has the same active site as ACE. If chitosan can block the binding of SARS-CoV-2 to ACE2 and remain the antihypertensive activity of ACE2, this study will have great scientific significance. Due to its unique biochemical and physicochemical properties, chitosan has been widely applied in the food and cosmetic industries as well as in the biomedical field. Chitosan occurs in three distinct crystalline polymorphic forms: α-, β- and γ-chitosan^14-16^. Chitosan is normally dissolved in acidic conditions and precipitates when the pH of the solution is above 7, which has greatly limited its utilization in cell and animal studies. Compared with α-chitosan, β-chitosan has weaker intramolecular and intermolecular hydrogen bonding forces, higher solubility, and greater biological activity.

Previous work indicated that β-chitosan (**Mendel®**) can be stably dissolved in a buffer solution with a similar pH and ionic concentration (pH 7.4, 20 mM) to human bodily fluids, as shown in **Extended Data Fig. 1**. Therefore, β-chitosan treatment was utilized to investigate its ability to block the interaction between SARS-CoV-2 S-RBD and ACE2 at the molecular, cellular, and animal levels. This work provides a promising approach to the prevention and control of SARS-CoV-2 infection using β-chitosan.

**Fig. 1.**
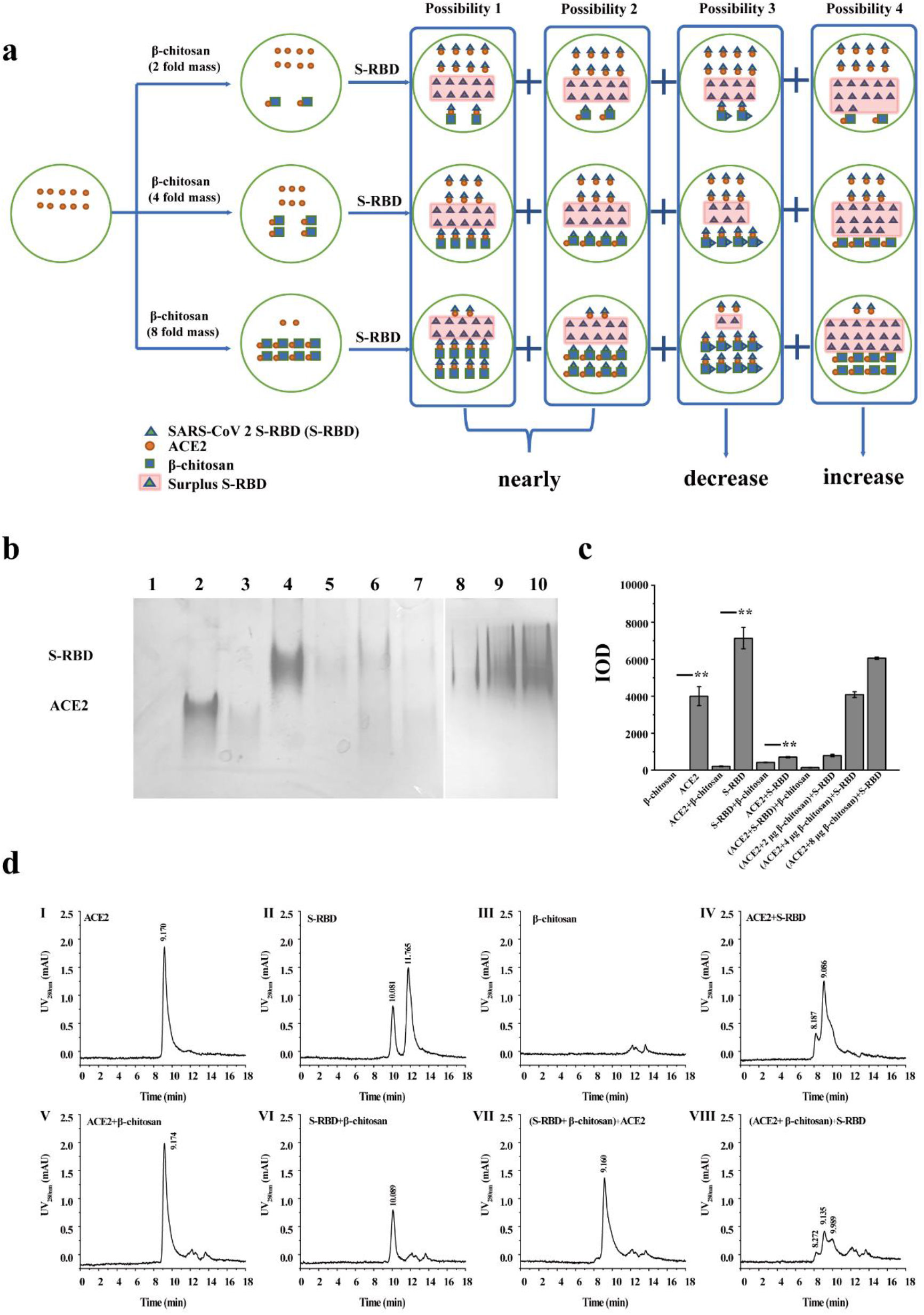
Binding interaction between β-chitosan and SARS-CoV-2 S-RBD/ACE2 analyzed via Native-PAGE and HPLC. **a**: These experiments were conducted to determine whether the conjugate of β-chitosan and ACE2 can bind to SARS-CoV-2 S-RBD (S-RBD). For the condition of excess ACE2 relative to β-chitosan, 1 μg of ACE2 was mixed and incubated with 2 μg, 4 μg, or 8 μg of β-chitosan for 20 min at 37°C, respectively. Then, 1 μg of S-RBD was added, followed by another 20 min co-incubation. Finally, Native-PAGE was used to analyze the samples. **b**: Native-PAGE analysis of the collected samples. (1). β-chitosan; (2). ACE2; (3). ACE2 + β-chitosan; (4). S-RBD; (5). S-RBD + β-chitosan; (6). ACE2 + S-RBD; (7). (ACE2 + S-RBD) + β-chitosan; (8). (ACE2 + 2 μg β-chitosan) + S-RBD; (9). (ACE2 + 4 μg β-chitosan) + S-RBD; (10). (ACE2 + 8 μg β-chitosan) + S-RBD. Samples (1-7) were obtained according to the procedures described in the Methods Section. Samples (8-10) were acquired using the method described for Fig. 1a. **c**: Integrated option density (IOD) was analyzed using Image J software. According to a two-tailed Student’s t-tests, the IODs of ACE2 and S-RBD incubated with β-chitosan (lane 3 and 5) diminished significantly compared with those of ACE2 and S-RBD (lanes 2 and 4). The grey value of the co-incubation mixture of ACE2 with S-RBD (lane 6) was diminished significantly compared to that of ACE2 or S-RBD, respectively. **d**: HPLC analysis of the collected samples. All samples were mixed and incubated for 20 min at 37°C. However, samples (7) and (8) underwent addition of β-chitosan or S-RBD and another 20 min incubation at 37°C, based on samples (4) and (5), respectively. (1). ACE2; (2). S-RBD; (3). β-chitosan; (4) ACE2 + S-RBD; (5). ACE2 + β-chitosan; (6). S-RBD + β-chitosan; (7). (S-RBD + β-chitosan) + ACE2; (8). (ACE2 + β-chitosan) + S-RBD.

## RESULTS

### I *In vitro* molecular interaction model for studying the effect of β-chitosan on the binding of the SARS-CoV-2 S-RBD with ACE2

An *in vitro* molecular interaction model was used to explore the effect of β-chitosan treatment on the binding of the SARS-CoV-2 S-RBD to ACE2. The co-incubation mixtures collected were analyzed by Native-PAGE. As shown in **Fig. 1b and Fig. 1c**, the IODs of ACE2 and SARS-CoV-2 S-RBD incubated with β-chitosan (lanes 3 and 5) decreased by 95.8% and 94.8% when compared with those of ACE2 and SARS-CoV-2 S-RBD (lane 2 and 4), respectively. This finding indicates that β-chitosan has a strong binding affinity to ACE2 and the SARS-CoV-2 S-RBD under normal physiological conditions *in vitro*. In addition, the IOD of the co-incubation mixture of ACE2 with SARS-CoV-2 S-RBD in lane 6 was reduced significantly compared with that of ACE2 or SARS-CoV-2 S-RBD, respectively, suggesting that SARS-CoV-2 S-RBD is strongly bound to ACE2. With the addition of β-chitosan, the IOD in lane 7 was decreased significantly, which demonstrated that β-chitosan treatment inhibited the binding of ACE2 and the SARS-CoV-2 S-RBD.

In order to confirm whether the conjugate of β-chitosan and ACE2 (termed β-chitosan∼ACE2) could rebind the SARS-CoV-2 S-RBD, we designed a creative experiment shown in **Fig. 1a**. Under the experimental condition of excess ACE2 relative to β-chitosan, the same amount of ACE2 was mixed and incubated with different amounts of β-chitosan. Then, this experiment was repeated with excess SARS-CoV-2 S-RBD added during the co-incubation. The samples were then analyzed by Native-PAGE. With increasing β-chitosan levels, β-chitosan∼ACE2 levels correspondingly increased and the surplus unbound ACE2 levels decreased. As a specific receptor binding domain of ACE2, the excess unbound SARS-CoV-2 S-RBD should bind to the excess ACE2. When the same amounts of SARS-CoV-2 S-RBD were added, it was unclear whether the final remaining SARS-CoV-2 S-RBD would bind with ACE2 and/or β-chitosan to form a conjugate. The following three situations could occur for the remaining SARS-CoV-2 S-RBD (**Fig. 1a**): (1) only ∼β-chitosan or ∼ACE2 could continually bind with the remaining SARS-CoV-2 S-RBD, then eventually the remaining SARS-CoV-2 S-RBD amounts would be the same and the IODs would show no difference; (2) both ∼β-chitosan and ∼ACE2 could continually bind with the remaining SARS-CoV-2 S-RBD, then the remaining SARS-CoV-2 S-RBD levels and IODs would gradually decrease; (3) neither ∼β-chitosan nor ∼ACE2 could continually bind with the surplus SARS-CoV-2 S-RBD, the remaining SARS-CoV-2 S-RBD and grey values would gradually increase. Native-PAGE results showed that the IODs for the SARS-CoV-2 S-RBD increased gradually (lane 8-10 in **Fig. 1c**), which is in accordance with the third speculation. These findings indicates that β-chitosan can bind with ACE2 and potently block the binding of the SARS-CoV-2 S-RBD to ACE2.

HPLC was also used to analyze the co-incubation mixtures collected. As shown in **Fig. 1d**, for ACE2 there was only a characteristic peak at 9.17 min (**Fig. 1d-**□), and two peaks were observed for SARS-CoV-2 S-RBD at 10.081 min and 11.765 min (**Fig. 1d-**□). This double peak is likely because the SARS-CoV-2 S-RBD is composed of polymers and monomers. Moreover, no peak for β-chitosan was observed in the current detection system (**Fig. 1d-**□). Two peaks were observed for the conjugate of the SARS-CoV-2 S-RBD and ACE2 (**Fig. 1d-**□), and the retention times (t_R_) shifted to 8.187 min and 9.086 min, respectively, indicating that the combination of SARS-CoV-2 S-RBD and ACE2 causes a shift in t_R_. For the co-incubation mixture of β-chitosan and ACE2, only one peak (9.17 min in **Fig. 1d-**□) was observed and the peak area was similar to that of ACE2 by itself, as shown in **Fig. 1d-**□. This finding suggested that the conjugate of β-chitosan and ACE2 present at atmospheric pressure dissociated under conditions of high pressure (40 MPa), which indicated that the binding interaction between β-chitosan and ACE2 is relatively weak. Interestingly, the conjugate of β-chitosan and the SARS-CoV-2 S-RBD showed a peak at 10.81 min with a peak area similar to the SARS-CoV-2 S-RBD in **Fig. 1d-**□, while the second peak disappeared (**Fig. 1d-**□). This finding illustrated that the binding interaction between β-chitosan and the SARS-CoV-2 S-RBD is very strong. Similar experiments were performed to verify whether the conjugate of β-chitosan and the SARS-CoV-2 S-RBD can bind with ACE2. As shown in **Fig. 1d-**□, only one ACE2 peak was observed in the mixture and the peak area was similar to that of ACE2 alone (**Fig. 1d-**□). This result revealed that the conjugate of β-chitosan and the SARS-CoV-2 S-RBD could not bind with ACE2, which was consistent with the above Native-PAGE analysis. If β-chitosan∼ACE2 dissociates at 40 MPa, the dissociated ACE2 would then bind with the SARS-CoV-2 S-RBD, leading to results similar to those in **Fig. 1d-**□. However, the results in **Fig. 1d-**□ revealed a significant difference, indicating that β-chitosan may interfere with the binding of the SARS-CoV-2 S-RBD to ACE2.

Therefore, the results of the *in vitro* molecular model indicate that β-chitosan can neutralize the SARS-CoV-2 S-RBD and block the binding of SARS-CoV-2 S-RBD to ACE2, suggesting that β-chitosan can shield ACE2 from being bound by the SARS-CoV-2 S-RBD.

### II *In vitro* Vero E6 cell model for studying the effect of β-chitosan treatment on the binding of the SARS-CoV-2 S-RBD with ACE2

Immunofluorescence (IF), flow cytometry (FCM) and Western blot (WB) analysis were used to investigate β-chitosan-mediated inhibition of the binding of the SARS-CoV-2 S-RBD to ACE2 in a Vero E6 cell model. Based on previous work that indicated treatment with β-chitosan could inhibit the expression of inflammation-related proteins in Vero E6 cells (**Extended Data Fig. 2**), a 500 μg/mL dose of β-chitosan was selected for further experiments.

**Fig. 2.**
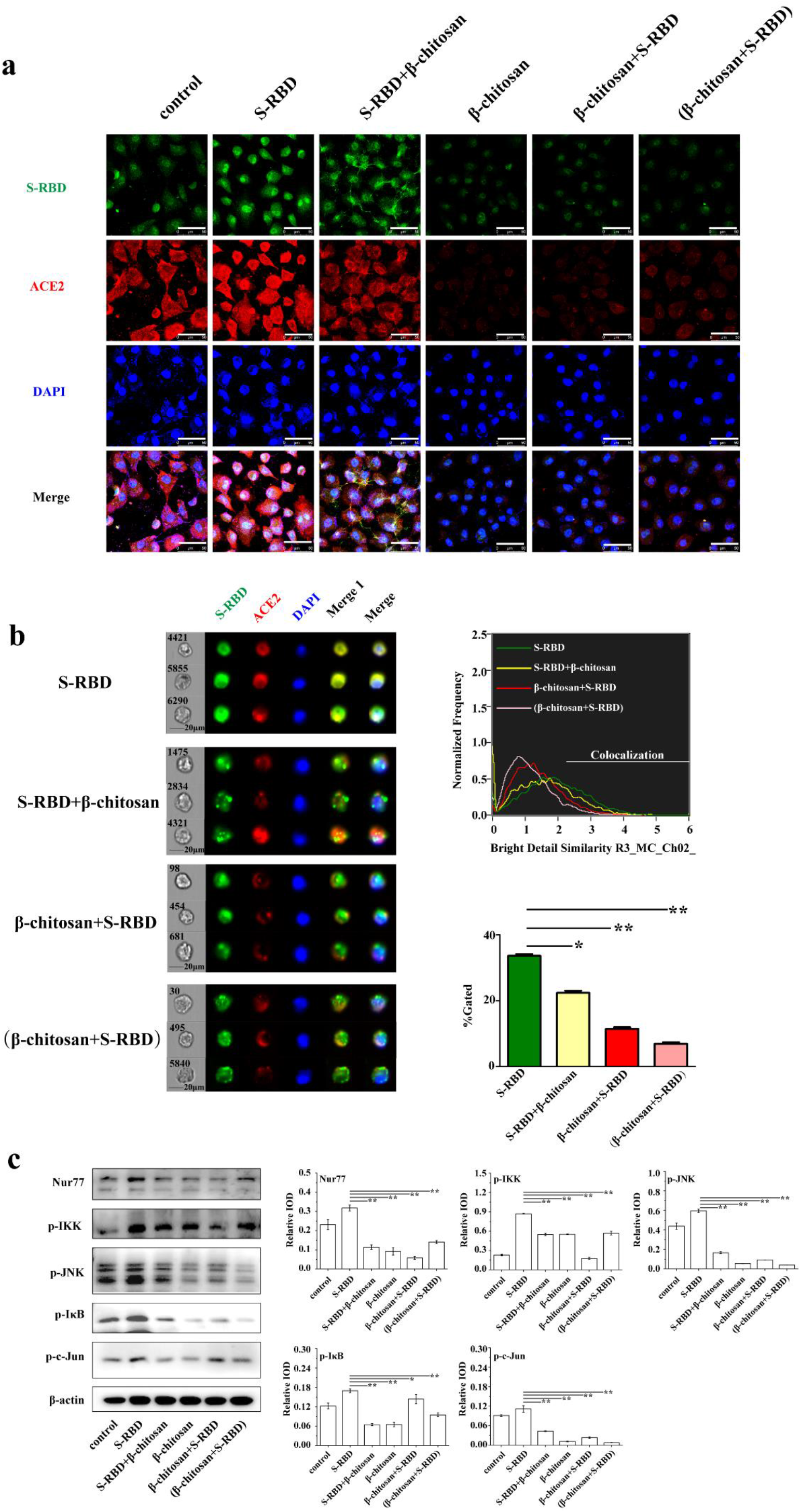
Effect of β-chitosan on binding of SARS-CoV-2 S-RBD to ACE2 assessed using a Vero E6 cell model. **a**: Effect of β-chitosan on binding of SARS-CoV-2 S-RBD (termed S-RBD in all the figures) to ACE2 detected via immunofluorescence. Single channel images of S-RBD, ACE2 and DAPI are shown. Moreover, a single overlapped image (the merge image) was merged and generated using the single channel images for S-RBD, ACE2, and DAPI. **b**: Effect of β-chitosan on S-RBD binding to ACE2 as determined using flow cytometry. Merge 1 was generated by merging the single channel images of S-RBD and ACE2. **c**: Western blot analysis of inflammation-related proteins. Cells were treated with S-RBD and/or β-chitosan as shown in Table 1. Relative IOD = IOD/β-actin. Scale bars, 50 μm (white).

IF analysis showed that ACE2 (red) and the SARS-CoV-2 S-RBD (green) presented strong fluorescence signals following the addition of the SARS-CoV-2 S-RBD into Vero E6 cells (**Fig. 2a-**□). Moreover, significant colocalization of ACE2 with SARS-CoV-2 S-RBD was observed, implying that the SARS-CoV-2 S-RBD binds to ACE2. When β-chitosan was added to the above system 10 min later, the colocalization of ACE2 and SARS-CoV-2 S-RBD was reduced (**Fig. 2A-**□), indicating that β-chitosan can prevent the binding of SARS-CoV-2 S-RBD to ACE2 exposed on the surface of Vero E6 cells. When β-chitosan and the SARS-CoV-2 S-RBD were added successively, the fluorescence intensities of ACE2 and the SARS-CoV-2 S-RBD were also significantly reduced (**Fig. 2A-**□). This suggested that β-chitosan treatment down-regulates the expression of ACE2 in Vero E6 cells, resulting in a decreased binding probability for the SARS-CoV-2 S-RBD with Vero E6 cells. When mixtures of β-chitosan and SARS-CoV-2 S-RBD were added to Vero E6 cells, the fluorescence intensities of ACE2 and SARS-CoV-2 S-RBD decreased significantly and no colocalization was observed (**Fig. 2A-**□), indicating that β-chitosan demonstrates a neutralizing activity against the SARS-CoV-2 S-RBD. In addition, it was also observed that the addition of β-chitosan alone significantly decreased ACE2 expression in Vero E6 cells (**Fig. 2A-**□).

**Table 1.**
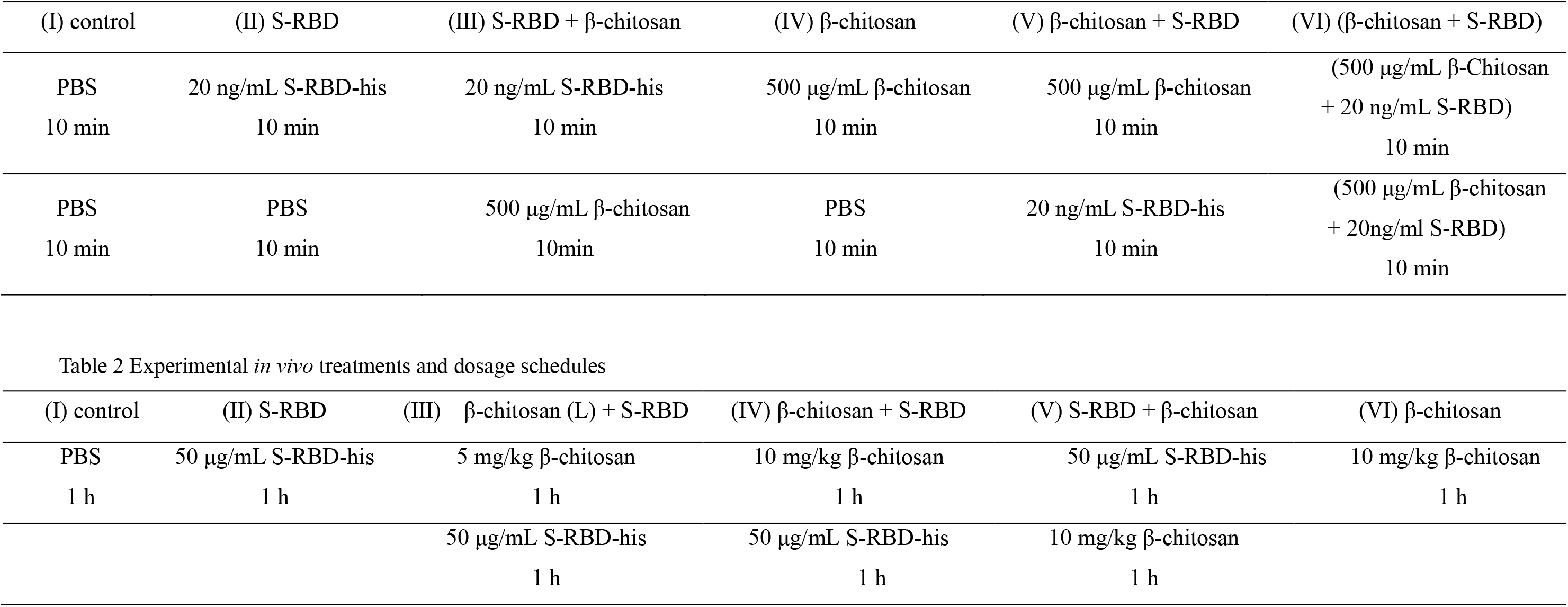
Experimental *in vitro* treatments and dosage schedules

FCM was used to detect the effect of β-chitosan treatment on Vero E6 cells infected with the SARS-CoV-2 S-RBD (**Fig. 2b**). Under the same parameters, 8000 cells were selected for analysis, and the colocalization of each experimental group was compared. It was found that the colocalization of ACE2 and the SARS-CoV-2 S-RBD reached 33.6% (green line in **Fig. 2b**) in Vero E6 cells following a 10 min incubation with the SARS-CoV-2 S-RBD. When β-chitosan was added into the above system 10 min later, the colocalization proportion fell to 22.4% (yellow line in **Fig. 2b**), indicating that the addition of β-chitosan decreased binding of the SARS-CoV-2 S-RBD to Vero E6 cells. When both β-chitosan and the SARS-CoV-2 S-RBD were added successively to Vero E6 cells in culture for 10 min (red line in **Fig. 2b**), respectively, the colocalization proportion was only 11.4%, indicating that β-chitosan could significantly diminish binding of the SARS-CoV-2 S-RBD with Vero E6 cells. When mixtures of β-chitosan and the SARS-CoV-2 S-RBD were added to Vero E6 cells, the colocalization proportion was only 6.91%, which further supports the conclusion that β-chitosan possesses neutralizing activity against the SARS-CoV-2 S-RBD.

The expression levels of proteins involved in inflammation response signaling pathways were analyzed by WB. p-JNK, p-IKK, p-IκB, and p-c-Jun are known to play a role in the response to inflammation. Nur77, an orphan member of the nuclear receptor superfamily, is expressed in macrophages following application of inflammatory stimuli^17^. The expression of Nur77, p-JNK, p-IKK, p-IκB, and p-c-Jun were all up-regulated significantly in Vero E6 cells following exposure to the SARS-CoV-2 S-RBD for 10 min (**Fig. 2c**). However, the expression of these proteins was significantly down-regulated when Vero E6 cells exposed to the SARS-CoV-2 S-RBD were treated with β-chitosan for 10 min. These results indicated that the SARS-CoV-2 S-RBD can induce inflammation in cells. However, the addition of β-chitosan inhibited the activation of inflammatory response signaling pathways, thus exerting an anti-inflammatory effect in cells.

### III *In vivo* animal model for examining effect of β-chitosan treatment on the binding of SARS-CoV-2 S-RBD to ACE2

FITC tags are often used to observe β-chitosan metabolism *in vivo*. However, the dissociation of β-chitosan-FITC during metabolism can lead to a false positive. Thus, β-chitosan was delivered intranasally to mice and Calcofluor White (CFW) was then used to observe its metabolic distribution ^18^. After intranasal administration for 1 h, 2 h, and 4 h, the metabolism and distribution of β-chitosan in the lungs of mice was analyzed. The blue fluorescence signal from the β-chitosan dyed with CFW was clearly distributed throughout the lungs of the mice (**Fig. 3a**). These results indicated that intranasally administered β-chitosan spreads throughout the lungs and attaches to lung epithelial cells over time, confirming the utility of this delivery route for the further evaluation of the antiviral effects of β-chitosan in the lung.

**Fig. 3.**
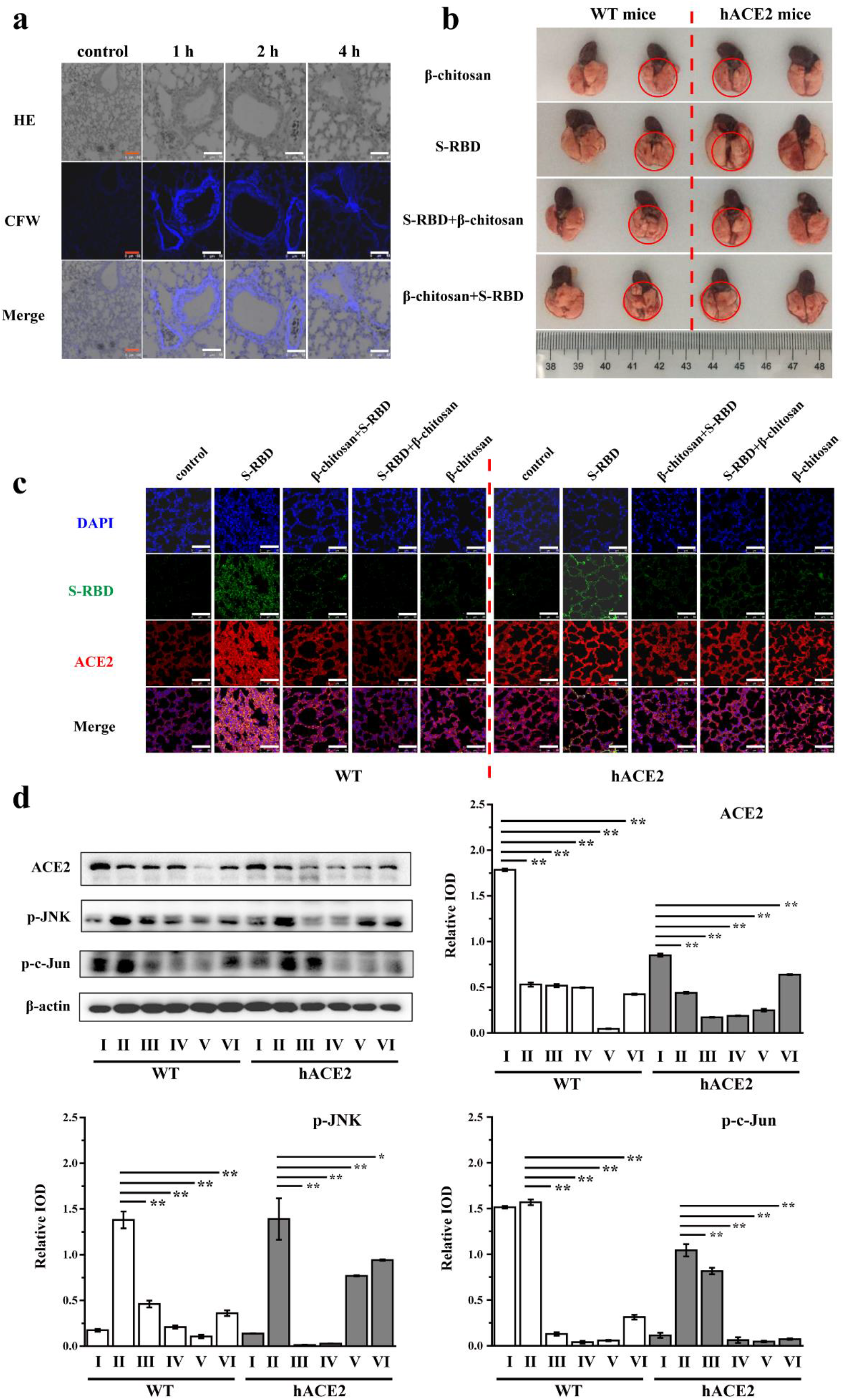
β-Chitosan distribution, histopathology, immunofluorescence, and effect on inflammation-related proteins in the lungs of WT and hACE2 mice. **a**: The distribution of β-chitosan was detected using CFW in the lungs of WT mice treated with 10 mg/kg β-chitosan for 1 h. WT and hACE2 mice were treated with S-RBD and/or β-chitosan as shown in Table 2. The merge image was generated by merging the single channel images of HE and CFW. **b**: Histopathology of lungs from WT and hACE2 mice. **c**: Colocalization of ACE2 with S-RBD as observed by immunofluorescence. **d**: Expression levels of ACE2 and inflammation-related proteins were analyzed by Western blot. Scale bars, 100 μm (red), 50 μm (white).

The preventive and alleviative effect of β-chitosan on the morphological lesion induced by SARS-CoV-2 S-RBD in mouse lungs was further observed (**Fig. 3b**). The lungs of hACE2 mice infected with the SARS-CoV-2 S-RBD were obviously swollen. However, the lungs of hACE2 mice treated with β-chitosan showed no difference from the control mice, indicating that β-chitosan treatment could inhibit the pathological changes in the lungs of hACE2 mice caused by the SARS-CoV-2 S-RBD. However, the SARS-CoV-2 S-RBD did not cause lung enlargement in wild-type (WT) mice, which indicated that the SARS-CoV-2 S-RBD has a specific effect on the morphological lesions present in hACE2 mice.

The expression of ACE2 and its colocalization with SARS-CoV-2 S-RBD for WT and hACE2 mice were determined using IF. In WT mouse lungs, the number of cells per unit area and the fluorescence intensity of ACE2 increased significantly in the SARS-CoV-2 S-RBD infected group as shown in **Fig. 3c**. However, the number of cells in hACE2 mouse lungs was significantly decreased and no obvious ACE2 immunofluorescence was observed in the SARS-CoV-2 S-RBD infected group. There were some similar phenomena observed between the two genotypes of mice. A visible immunofluorescence signal for the SARS-CoV-2 S-RBD and the colocalization of the SARS-CoV-2 S-RBD with ACE2 could be observed, implying that the lung cells of both WT and hACE2 mice can bind with the SARS-CoV-2 S-RBD through their ACE2 receptor. For the prevention groups (treated with β-chitosan and SARS-CoV-2 S-RBD successively) and treatment groups (treated with SARS-CoV-2 S-RBD and β-chitosan successively), no obvious SARS-CoV-2 S-RBD immunofluorescence signal was observed, which indicated that β-chitosan has a therapeutic effect by blocking the binding of SARS-CoV-2 S-RBD to lung cells in both types of mice.

In addition, the expression of ACE2 and inflammation-related proteins were analyzed and compared by WB. For the groups infected with the SARS-CoV-2 S-RBD (SARS-CoV-2 S-RBD group), ACE2 expression was distinctly reduced in comparison with the uninfected control groups of WT and hACE2 mice, respectively. For the prevention groups and treatment groups, the expression of ACE2 in both two types of mice was significantly decreased compared with that of the SARS-CoV-2 S-RBD groups (**Fig. 3d**). Moreover, expression of inflammation-related proteins (such as p-JNK and p-c-Jun) in the SARS-CoV-2 S-RBD groups was significantly up-regulated when compared with the control group (**Fig. 3d**). However, the expression levels of these proteins in the experimental groups treated with β-chitosan were significantly down-regulated. The results indicated that β-chitosan treatment could modulate ACE2 expression to decrease SARS-CoV-2 S-RBD infection. While the presence of the SARS-CoV-2 S-RBD promotes inflammation in the lung tissue, β-chitosan treatment can inhibit the activation of inflammation-related signaling pathways and exerts an anti-inflammatory effect. In general, the results of the animal model studies are consistent with those of the cell model.

### IV Regulation of ACE2 expression by β-chitosan

The above results suggested that β-chitosan treatment could significantly down-regulate ACE2 expressions in Vero E6 cells and lung cells in hACE2 mice. However, this decrease in ACE2 expression did not cause inflammation, on the contrary, β-chitosan treatment appeared to inhibit the inflammation induced by the SARS-CoV-2 S-RBD. The above mentioned results indicated that β-chitosan could reduce ACE2 expression without affecting its antihypertensive and anti-inflammatory effects, which implies that there might be another mechanism behind the decreased expression of ACE2. It was reported that ACE2 could be cleaved by the ADAM17 metalloproteinase, preventing the binding of SARS to the ACE2 receptor^19^. However, whether the decreased ACE2 expression observed in this study is related to ADAM17 activity is unknown. Therefore, an ADAM17 inhibitor was used in the context of IF and WB analyses to determine the role of this activity in this system.

The results of these experiments demonstrated that β-chitosan treatment significantly decreased the expression of ACE2 in Vero E6 cells (**Fig. 4**). When the ADAM17 inhibitor (TAPI-1) was added in advance, the expression of ACE2 was significantly up-regulated compared with that of the no TAPI-1 treatment group. Thus, these findings support the conclusion that the decreased ACE2 expression observed with β-chitosan treatment is caused by the activation of ADAM17. Moreover, the activation of ADAM17 could improve the cleavage of the ACE2 extracellular juxtamembrane region, which possesses catalytic activity against Ang II. The release of the catalytically active ectodomain into the extracellular environment can result in decreased binding interactions between cells and the SARS-CoV-2 S-RBD, thus giving the appearance of down-regulated ACE2 expression. Furthermore, the cleaved extracellular domain of ACE2 retains its catalytic activity that degrades Ang II and promotes antihypertensive effects.

**Fig. 4.**
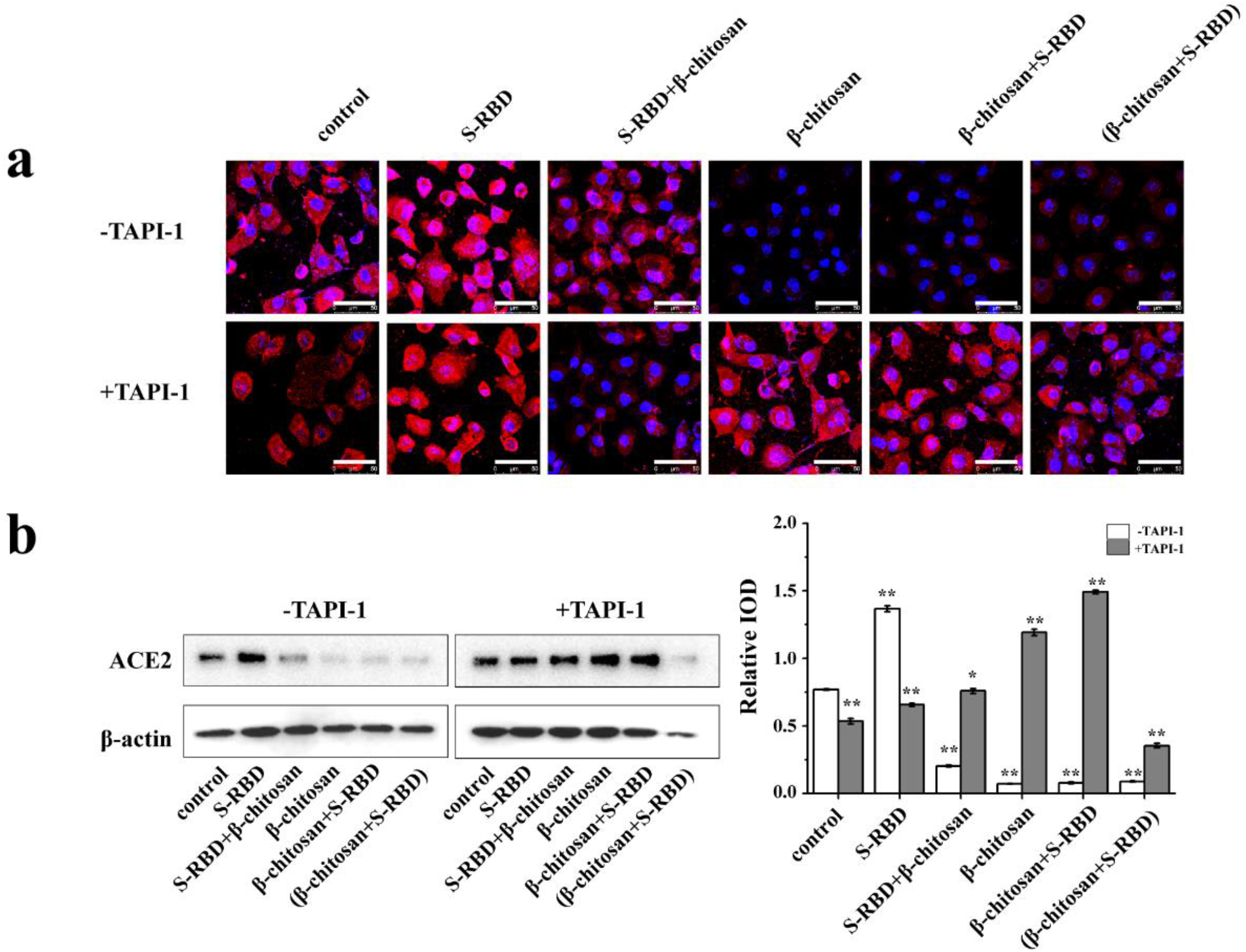
Verification of β-chitosan-mediated activation of ADAM17 cleavage of ACE2. **a**: Effect of ADAM17 inhibitor TAPI on the expression of ACE2 as detected by immunofluorescence. Cells were treated with 8 μM of an ADAM17 inhibitor TAPI or with PBS for 30 min, and the cells were then treated as shown in Table 1. Scale bars, 50 μm (white). **b**: Expression levels of ACE2 and inflammation-related proteins were analyzed by Western blot. All groups were compared with the no TAPI-1 control group. Scale bars, 50 μm (white).

## DISCUSSION

In this work, binding of SARS-CoV-2 S-RBD to β-chitosan was confirmed by Native-PAGE and HPLC. Interestingly, β-chitosan can bind with high affinity to the SARS-CoV-2 S-RBD without dissociation under high pressure. Thus, an additional *in vitro* molecular experiment was designed to verify that the conjugate of β-chitosan and ACE2 could no longer bind with the SARS-CoV-2 S-RBD. IF studies revealed that the SARS-CoV-2 S-RBD interacted with β-chitosan extracellularly, preventing it from binding to ACE2 on the surface of Vero E6 cells. Therefore, β-chitosan plays an important role similar to that of an antibody, neutralizing the SARS-CoV-2 S-RBD and effectively blocking the binding of SARS-CoV-2 S-RBD to ACE2 and preventing infection of the cells.

In the human body, ACE is an angiotensin converting enzyme that catalyzes the conversion of angiotensin I (Ang I) to Ang □. Ang □ plays a crucial role in contracting blood vessels and promoting inflammation through the angiotensin type 1 receptor (AT1R). However, ACE2 plays a distinct role from ACE in that it can catalyze the conversion of Ang □ to vasodilator angiotensin 1-7 (Ang 1-7), which can in turn dilate blood vessels and diminish inflammation through the Mas receptor. Thus, it could be hypothesized that a system formed on the basis of the ACE-Ang □-AT1R and ACE2-Ang 1-7-Mas axes maintains the blood pressure homeostasis in the human body^20,21^. Due to the endocytosis induced by the binding of SARS-CoV-2 with ACE2, ACE2 on the surface of cells is almost entirely consumed, causing Ang □ levels to increase significantly, leading to a cytokine storm and multiple organ damage^7,22^. Therefore, once an individual becomes infected by SARS-CoV-2, whether ACE2 is beneficial or detrimental to the individual has become a hotly debated scientific issue. Acute respiratory distress syndrome (ARDS) is one of the leading causes of death for patients infected with COVID-19 and SARS. As a result, the relationship between ARDS and ACE2 has attracted significant attention. Imai ^23^ found that ARDS lesions in mice with the ACE2 gene knocked out were significantly more severe than those in the control group, while the application of recombinant ACE2 could significantly reduce the occurrence of ARDS in mice, suggesting that ACE2 expression has an obvious protective effect with regard to ARDS in mice. Kuba et al. ^24^ reported that the expression of ACE2 was down-regulated *in vivo* and *in vitro* in the presence of recombinant SARS-CoV-S protein, resulting in pulmonary edema, deterioration in lung function, and ARDS in mice. It has been also reported that ACE2 expression in the myocardium of mice and patients infected with SARS-CoV was decreased significantly^7^. These findings suggest that ACE2 expression levels are closely related to the severity of SARS lesions. Liu et al. ^25^ found that Ang □ levels in the plasma of COVID-19 patients were significantly higher than those of healthy controls, and moreover, Ang □ levels were linearly related to virus titer and the degree of lung injury, indirectly indicating that those phenomena were all induced by decreased ACE2 expression.

Native-PAGE analysis demonstrated that ACE2 can be bound by β-chitosan, which prevents the binding of the SARS-CoV-2 S-RBD. The IF and FCM results from the cell and animal models revealed that β-chitosan treatment had a significant effect on preventing SARS-CoV-2 infection. WB analyses showed that the expression levels of ACE2 in Vero E6 cells and hACE2 mice infected with the SARS-CoV-2 S-RBD were significantly decreased and inflammation responses were also activated. Compared to the individual SARS-CoV-2 S-RBD group, the expression of ACE2 in each group of cells or tissues treated with β-chitosan was significantly decreased, which in turn decreased the inflammatory response driven by the presence of the SARS-CoV-2 S-RBD. This is an interesting and important discovery. Based on the known regulatory mechanisms related to ACE2 expression, ACE2 expression is down-regulated as a result of virus binding, which causes internalization and degradation of ACE2. Moreover, the transmembrane protease ADAM17 can cleave the juxtamembrane region of ACE2 and release the active extracellular domain, which can catalyze the degradation of Ang II in the extracellular environment^19,26^. Thus, this led to speculation with regard to whether there was a mechanism by which β-chitosan, ADAM17, and ACE2 interact to drive an anti-SARS-CoV-2 effect. Accordingly, an ADAM17 inhibitor (TAPI-1) was used to test this hypothesis, and the results indicated that the decreased ACE2 expression level observed in the experimental groups treated with β-chitosan was reversed upon addition of an ADAM17 inhibitor. Compared with the no inhibitor group, the expression of ACE2 was significantly increased (* *P* < 0.05; * * *P* < 0.01). This finding suggested that β-chitosan can activate ADAM17 to enhance the cleavage of the extracellular juxtamembrane region of ACE2, releasing the extracellular domain with catalytic activity into the extracellular environment, thus reducing the binding and internalization of SARS-CoV-2 S-RBD to susceptible cells. Moreover, the cleaved extracellular region of ACE2 retains its catalytic activity and continues to degrade Ang II and prevent inflammation. This discovery that β-chitosan treatment can prevent the binding of the SARS-CoV-2 S-RBD to ACE2 is shown in **Fig. 5**. Therefore, ADAM17 is a most likely relevant target for the treatment of SARS-CoV-2.

**Fig. 5.**
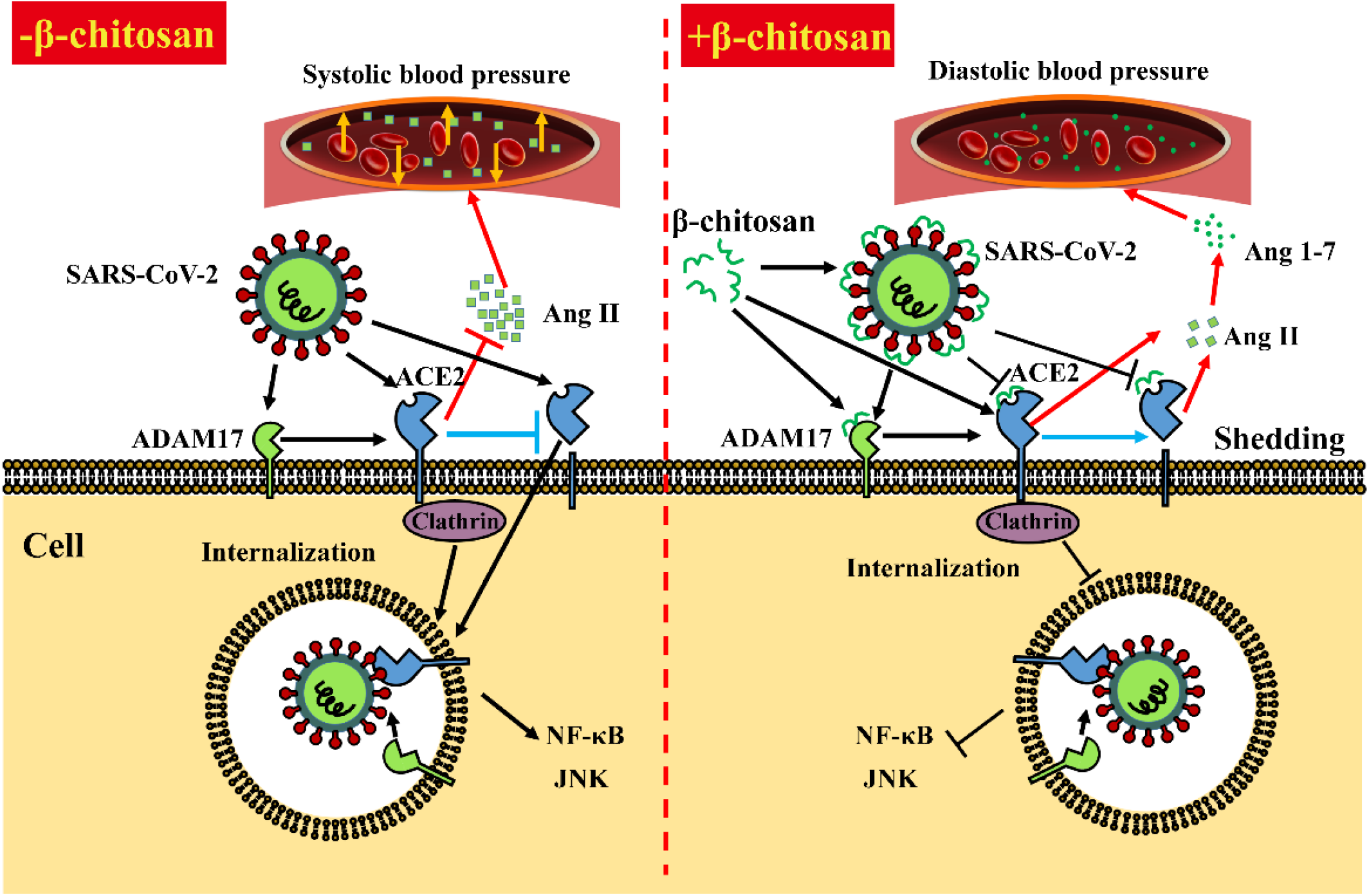
Mechanism for β-chitosan-mediated inhibition of SARS-CoV-2 S-RBD/ACE2 binding. -β-chitosan: control group for testing SARS-CoV-2 S-RBD/ACE2 binding. SARS-CoV-2 gains entry into host cells via the binding and internalization of ACE2 in a clathrin-dependent manner. Thus, the levels of the extracellular juxtamembrane region of ACE2, which can catalyze the degradation of Ang II, are decreased, resulting in increased Ang II levels and elevated blood pressure^19^. Moreover, ADAM17 cleaves few extracellular domain of ACE2. Meanwhile, the NF-κB and JNK signaling pathways are activated along with some related-inflammatory pathways. The arrowheads indicate the proposed process. The “T” shaped arrows represent an inhibitory effect. **+β-chitosan: treatment group for testing SARS-CoV-2 S-RBD/ACE2 binding**. In the presence of β-chitosan, SARS-CoV-2 S-RBD and ACE2 are bound and shielded by β-chitosan, which blocks the binding and subsequent internalization of SARS-CoV-2 S-RBD to ACE2. ADAM17 is activated by β-chitosan to enhance the cleavage of the extracellular domain of ACE2. The active ectodomain of ACE2 remains catalytically active and degrades Ang II, thus promoting maintenance of a stable blood pressure. In addition, the NF-κB and JNK signaling pathways are also inhibited.

Recently, Waradon Sungnak reported^27^ that ACE2 and TMPRSS2 were highly expressed in nasal goblet cells and ciliated cells that produce mucus. Moreover, the virus concentration in nasal swabs from respiratory disease patients infected by SARS-CoV-2 was higher than that of pharyngeal swabs, suggesting that nasal passages may be a channel for initial infection and transmission of the virus. By administering β-chitosan intranasally, we found that it can be effectively distributed in lung tissue. This result provides a precondition for β-chitosan to block the binding of SARS-CoV-2 S-RBD to ACE2 *in vivo*. Furthermore, β-chitosan which can be safely degraded, displays good biocompatibility in the brain tissue of rats ^28^. Kean also found that β-chitosan exhibited no toxicity in acute toxicity tests performed with mice and no irritation to the eyes and skin of rabbits^29^. Thus, delivery of β-chitosan by intravenous injection appears to be safe for humans. In this study, the infection capacity of the SARS-CoV-2 S-RBD and the therapeutic effect of β-chitosan administration were compared using WT and hACE2 mice. The results of these experiments suggested that the SARS-CoV-2 S-RBD could induce severe inflammation throughout the lungs of hACE2 mice, but there was no similar pathological phenomenon observed in WT mice, which is consistent with the results of Bao’s work^30^. However, immunofluorescence analysis of tissue samples showed that both lung epithelial cells from both WT and hACE2 mice could bind with SARS-CoV-2 S-RBD. The results therefore indicated that the SARS-CoV-2 S-RBD has different pathogenic effects on WT mice and hACE2 mice, with especially high pathogenicity in the hACE2 host, suggesting that the hACE2 mouse model is an indispensable scientific tool for screening drugs against SARS-CoV-2.

In this work, the inhibitory effect of β-chitosan on the binding of the SARS-CoV-2 S-RBD to ACE2 was investigated at three levels (molecular, cellular and animal). The results revealed that β-chitosan treatment can potently block the binding of the SARS-CoV-2 S-RBD to ACE2 and inhibit the occurrence of inflammation. This study presents a promising and important research with a great value and the prospect of application of β-chitosan as a therapy against SARS-CoV-2. With the current epidemic raging, it is time to share our research results in a timely manner and carry out additional studies to improve our collective understanding of this virus.

## Materials and methods

### Ethics statement

All procedures in this study involving animals were reviewed and approved by the Minnan Normal University Animal Ethics and Welfare Committee (MNNU-AEWC-2020001).

### Antibodies and reagents

Anti-His-Tag, anti-p-IKK, anti-p-JNK, and anti-p-c-Jun antibodies, FITC-conjugated donkey anti-mouse IgG, and Cy3-conjugated donkey anti-rabbit IgG were purchased from Affinity Biosciences, Inc. (Cincinnati, OH, USA). Anti-ACE2 antibody, HRP-conjugated anti-mouse IgG, and HRP-conjugated anti-rabbit IgG were obtained from Abcam (Cambridge, UK). Anti-β-actin antibody was purchased from Cell Signaling Technology (Beverly, MA, USA). CFW was provided by Shanghai Yuanye Bio-Technology Co., Ltd. (Shanghai, China). TAPI-1 was purchased from APExBIO Technology LLC (Houston, TX, USA). SARS-CoV-2 S-RBD and ACE2 proteins were obtained from Novoprotein Scientific Inc. (Shanghai, China).

Dulbecco’s modified Eagle medium (DMEM) was purchased from Invitrogen Corp. (Carlsbad, CA, USA). 10% fetal bovine serum (10% FBS) was purchased from Thermo Fisher Scientific Co., Ltd. (Waltham, MA, USA). β-Chitosan was provided by Mengdeer (Xiamen) Biotechnology Co., Ltd. (Xiamen, China).

Other chemicals and reagents used in this work were of analytical reagent grade.

### Binding of β-chitosan with ACE2 or SARS-CoV-2 S-RBD

In order to be compatible with the conditions required for HPLC analysis, proteins (SARS-CoV-2 S-RBD and ACE2) and β-chitosan were dissolved in a 0.02 M Tris-HCl buffer solution (pH 6.8) in advance. SARS-CoV-2 S-RBD or ACE2 was mixed with a suitable amount of β-chitosan and incubated at 37°C for 20 min. To test the effect of the presence of β-chitosan on the binding of ACE2 to the SARS-CoV-2 S-RBD, β-chitosan was added to the co-incubation mixture containing ACE2 and SARS-CoV-2 S-RBD and incubated together in a similar manner. The sample mixtures were then analyzed by Native-PAGE and SEC-HPLC. The amounts of proteins and β-chitosan used were determined based on the detection limit of Native PAGE and the relative molecular weights of the two proteins (SARS-CoV-2 S-RBD and ACE2).

### Cell experiments

Vero E6 cells (ATCC, CRL-1586) were cultured in DMEM supplemented with 10% (vol/vol) FBS, 50 units/mL penicillin, and 50 μg/mL streptomycin in a humidified atmosphere (5% CO_2_) at 37°C. Cells were grown to 80% confluency prior to undergoing any treatments for experiments.

8 μM TAPI-1 was added to Vero E6 cells 30 min in advance of any additional treatment conditions.

### Mouse experiments

Specific pathogen-free, 8-week-old male transgenic hACE2 mice and WT male mice (C57BL/6 background) were purchased from the experimental animal center of GemPharmatech (Nanjing, China). All mice were maintained in an animal room with a 12-h light/12-h dark cycle at the Laboratory Animal Center in Minnan Normal University (China). Transgenic mice were generated via microinjection of the mouse *ACE2* promoter, which delivers the human *ACE2* coding sequence to the pronuclei of fertilized ova from C57BL/6 mice. The integrated human ACE2 sequence was then confirmed using PCR as reported previously^31^. Human ACE2 is mainly expressed in the lung, heart, kidneys, and intestines of transgenic mice. β-Chitosan doses were intranasally administrated according to a previously published method^32^. SARS-CoV-2 S-RBD and/or β-chitosan were inoculated intranasally into hACE2 and WT mice at a dose of 10 mg/kg body weight, respectively, and an equal volume of PBS was used as control. Following treatment, mice were dissected to collect different tissues in order to observe any histopathological changes.

### Flow cytometry

When Vero E6 cells had grown to 80% confluency, they were treated according to the experimental design. Following treatment, the cells were harvested, washed with PBS and fixed overnight at 4°C with 200 μL of precooled paraformaldehyde (4%). Fixed cells were collected after centrifugation, washed twice with precooled PBS, sealed with 10% donkey serum for 15 min, and washed again with precooled PBS. Then, anti-His-Tag (1:200, Affinity, T0009, USA) and anti-ACE2 antibodies (1:100, Abcam, ab15348, USA) were added and the cells were incubated overnight at 4°C. Subsequently, the cells were washed three times with PBS, and FITC-conjugated donkey anti-mouse IgG (1:300, Affinity, USA), Cy3-conjugated donkey anti-rabbit IgG (1:300, Affinity, USA) and DAPI (1:2000, Affinity, USA) were added and incubated at 37°C for 60 min in the dark. The cells were then washed, re-suspended in precooled PBS and analyzed using flow cytometry (Flowsight, Merk Millipore, USA). The instrument was adjusted according to a previously reported procedure^33^. The colocalization of FITC-His-S-RBD and Cy3-ACE2 was analyzed using the colocalization analysis application Wizard in IDEAS software.

### CFW

Paraffin sections were dewaxed into water, washed with distilled water, and dried. CFW fluorescent staining solution was added and incubated in the dark for 10 min to dye the sections. Then, the sections were washed in distilled water and underwent routine HE staining. After washed with distilled water, the slides were subjected to gradient ethanol dehydration. Finally, the sections were cleared with xylene and sealed with neutral gum.

### Confocal microscopy

For immunofluorescence on cell culture, cells grown on glass cover-slips were fixed with 4% paraformaldehyde solution at 4°C overnight. Then permeabilized with 0.1 % triton X-100 for 10 min. The following primary antibodies His-Tag (1:200, Affinity, T0009, USA) and ACE2 (1:200, Abcam, ab15348, UK) were used. Then FITC-conjugated donkey anti-mouse IgG (1:400) or Cy3-conjugated donkey anti-rabbit IgG (1:400) was added at room temperature for 1 h.

For section immunofluorescence, fresh samples were isolated and fixed in 4% paraformaldehyde solution at 4°C overnight. After dehydration through gradient ethanol, samples were embedded in paraffin and sectioned. The following primary antibodies His-Tag (1:200, Affinity, T0009, USA) and ACE2 (1:200, Abcam, ab15348, UK) were used. Slides were subjected to standard immunohistochemistry staining.

### Native-PAGE analysis

A 6% resolving gel and a 5% condensing gel were used for Native-PAGE analysis. Protein solutions were diluted to a concentration of 0.5∼2.0 mg/mL. The amount of sample loaded was 2 µL of protein marker and 20 µL of each protein sample. The protein separation was accomplished by application of 100 V for 120 min. The gel was visualized by silver staining following electrophoresis according to a previously published method ^34^ with some modifications. The stained protein gel was scanned using a Gel Doc 2000 imaging system (Bio-Rad, Hercules, CA, USA). The protein signals were evaluated by integrated option density (IOD) using Image J software (National Institutes of Health, Bethesda, MD, USA).

### SEC-HPLC analysis

Analytical SEC-HPLC was performed using an Agilent 1200 HPLC system (Agilent Technologies, Santa Clara, CA, USA) with a Waters Ultrahydrogel^TM^ 1000 column (12 µm, 7.8 mm × 300 mm, Waters, Milford, MA, USA). The mobile phase of 0.2 M Tris-HCl buffer (pH 6.8) underwent 0.22 μm membrane filtration and degassing prior to use. The flow rate was 1 mL/min, and the sample injection volume was 20 µL. The chromatographic run was monitored online at 280 nm by a UV detector. The integrated peak areas of each component were used to evaluate the amount of protein present.

### Western blot analysis

Cells and mouse tissues were homogenized in cell lysis buffer. Proteins from cell or tissue lysates were separated using sodium dodecyl sulfate polyacrylamide gel electrophoresis (SDS-PAGE) and transferred to a nitrocellulose membrane. The membrane was then blocked in 5% non-fat milk for 1 h at room temperature and incubated with primary antibodies (including anti-ACE2, anti-p-IKK, anti-p-JNK, anti-p-c-Jun, and anti-β-actin antibodies) overnight at 4°C. The membrane was then washed with Tris buffered saline buffer with Tween 20 (TBST) for three times and incubated with secondary antibody (HRP-conjugated anti-IgG) for 1 h. The final immunoreactive products were observed using an Omega Lum C Gel Imaging system (Aplegen, USA), and the protein signals were evaluated by IOD using Image J software.

### Statistical analysis

All data were analyzed with GraphPad Prism 8.0 software. Two-tailed Student’s *t*-tests were applied for statistical analysis. For all data, a two-sided *P* value < 0.05 was considered statistically significant (\**P* < 0.05, \*\**P* < 0.01).

## Acknowledgements

This work was financially supported by the National Natural Science Foundation of China (81903665) and the Fujian Provincial Department of Finance (FJDF[2019]0926).

