## Supplementary Figures for "Effect of β-chitosan on the binding interaction between SARS-CoV-2 S-RBD and ACE2"

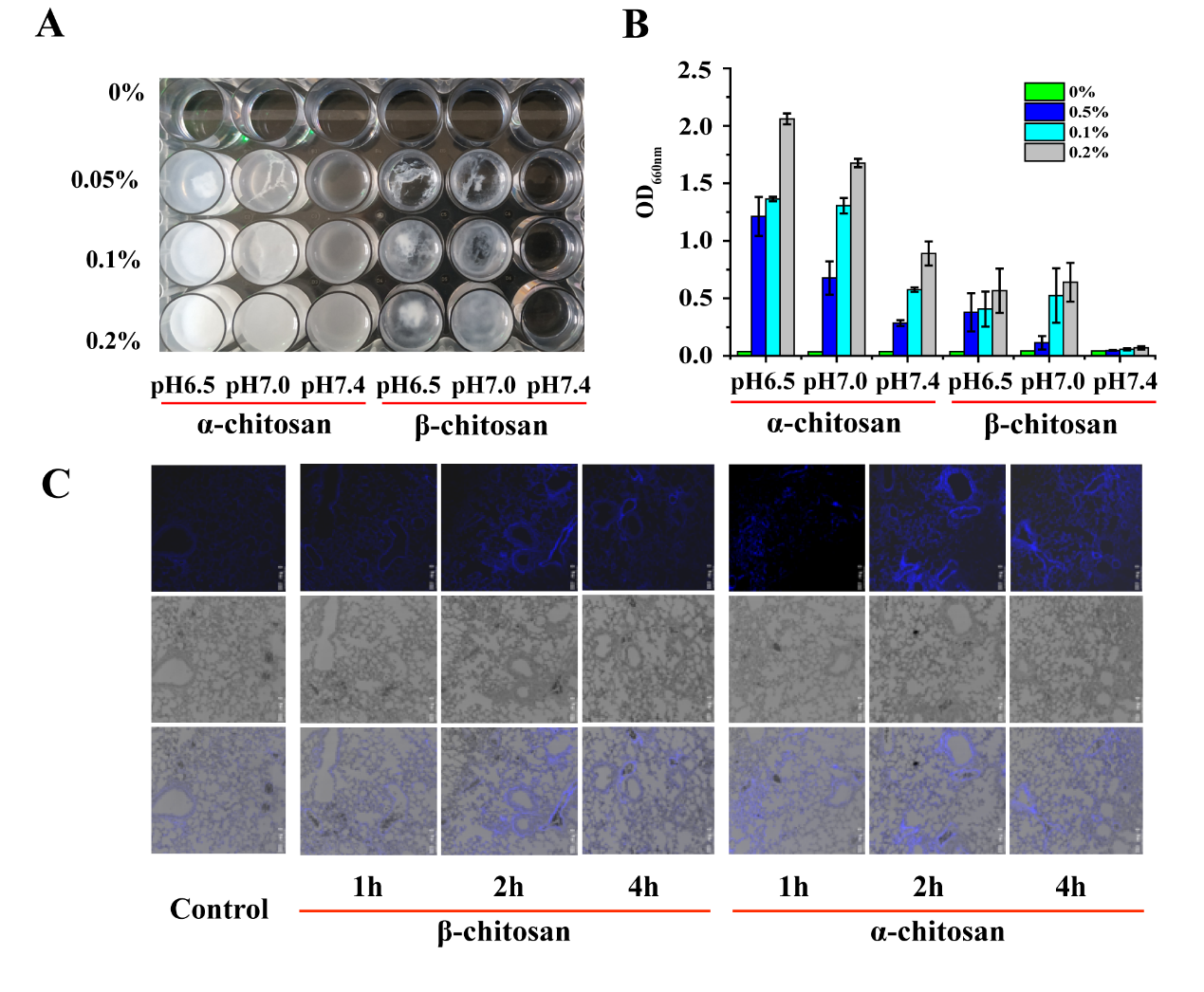

**Extended Data Fig. 1 Comparison of the solubility of α-chitosan and β-chitosan.** A: Chitosan solutions of different pHs (6.5-7.4) and different concentrations (0-0.2%) were prepared in 20 mM phosphate buffer saline solution. 30 min after preparing the solutions, turbidity was determined at 660 nm using a microplate reader (Infinite M200 Pro, Tecan Group Ltd., San Jose, USA). B: Graph comparing the turbidities of the α-chitosan and β-chitosan solutions under different conditions.

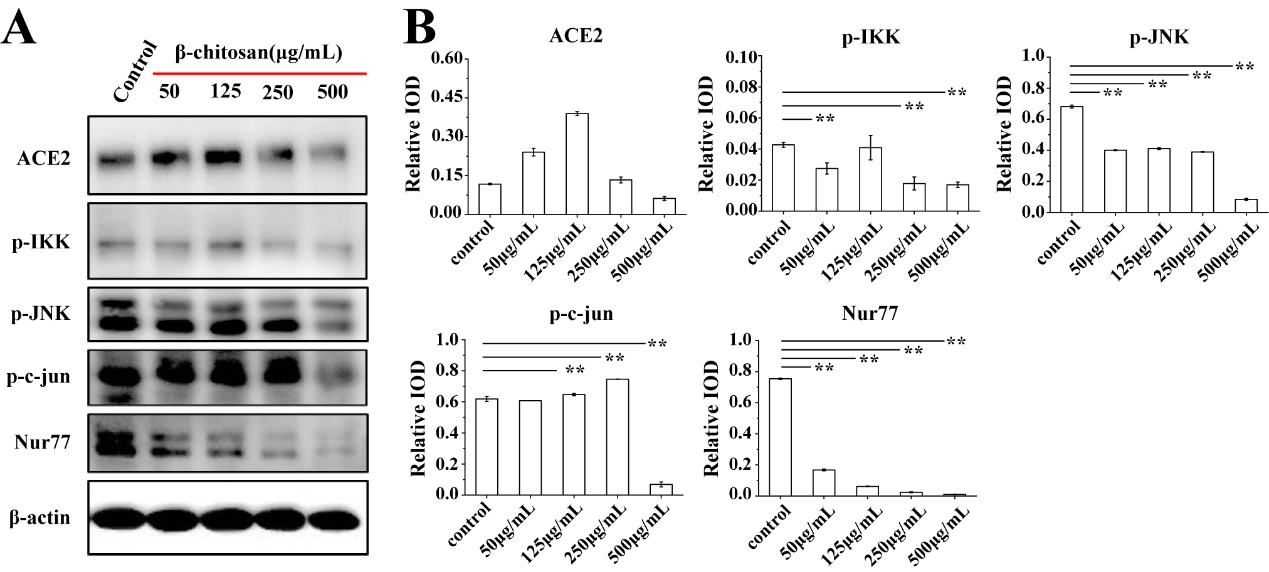

**Extended Data Fig. 2** Effect of β-chitosan dose on levels and activation of inflammation-related proteins in Vero E6 cells. A: WB analysis of inflammation-related proteins following treatment with different doses of β-chitosan (0-500 μg/mL). B: IODs for WBs of inflammation-related proteins. Relative IOD = IOD/β-actin.
